# Phylovar: Towards scalable phylogeny-aware inference of single-nucleotide variations from single-cell DNA sequencing data

**DOI:** 10.1101/2022.01.16.476509

**Authors:** Mohammadamin Edrisi, Monica V. Valecha, Sunkara B. V. Chowdary, Sergio Robledo, Huw A. Ogilvie, David Posada, Hamim Zafar, Luay Nakhleh

## Abstract

Single-nucleotide variants (SNVs) are the most common variations in the human genome. Recently developed methods for SNV detection from single-cell DNA sequencing (scDNAseq) data, such as SCIΦ and scVILP, leverage the evolutionary history of the cells to overcome the technical errors associated with single-cell sequencing protocols. Despite being accurate, these methods are not scalable to the extensive genomic breadth of single-cell whole-genome (scWGS) and whole-exome sequencing (scWES) data.

Here we report on a new scalable method, Phylovar, which extends the phylogeny-guided variant calling approach to sequencing datasets containing millions of loci. Through benchmarking on simulated datasets under different settings, we show that, Phylovar outperforms SCIΦ in terms of running time while being more accurate than Monovar (which is not phylogeny-aware) in terms of SNV detection. Furthermore, we applied Phylovar to two real biological datasets: an scWES triple-negative breast cancer data consisting of 32 cells and 3375 loci as well as an scWGS data of neuron cells from a normal human brain containing 16 cells and approximately 2.5 million loci. For the cancer data, Phylovar detected somatic SNVs with high or moderate functional impact that were also supported by bulk sequencing dataset and for the neuron dataset, Phylovar identified 5745 SNVs with non-synonymous effects some of which were associated with neurodegenerative diseases. We implemented Phylovar and made it publicly available at https://github.com/mae6/Phylovar.git.

## Introduction

With the advent of the first single-cell sequencing (SCS) techniques [20, 27], the fields of single-cell genomics, transcriptomics, proteomics, and epigenetics have witnessed remarkable growth over the last decade. Single-cell sequencing technologies have impacted our understanding in different fields of biology including developmental biology, immunology, microbiology, and cancer biology [30, 21, 16, 14, 28]. Single-cell DNA sequencing (scDNAseq), as one of the SCS technologies, provides insights into the somatic evolutionary process by sequencing the genomic contents of a complex tissue at a single-cell resolution [20, 21]. Preparing scDNAseq data requires a whole-genome amplification (WGA) process to amplify the DNA material of a single cell to suffice the amount of DNA needed for sequencing. [35, 14]. WGA technologies, such as multiple displacement amplification (MDA) [25, 3] and multiple annealing and looping-based amplification cycles (MALBAC) [37] can elevate the noise level in scDNAseq data. The scDNAseq technical errors include allelic dropout (ADO), false-positive (FP) errors, false-negative (FN) errors, and non-uniform coverage [21, 35]. ADO refers to cases where only one of the two alleles in a heterozygous mutation is amplified, resulting in the loss of the mutated allele. FP artifacts can appear due to uneven amplification or at the early stages of the amplification when the original nucleotide is substituted randomly. The non-uniform coverage over different genomic loci may result in missing data due to zero or insufficient coverage. The scDNAseq-specific technical errors fuelled the development of tools such as Monovar [36] and SCcaller [5] for detecting single-nucleotide variations (SNVs) from scDNAseq data. Although Monovar and SCcaller account for uneven coverage and scDNAseq-specific errors, more recent methods, SCIΦ [24] and scVILP [6], showed further improvement in overcoming the scDNAseq-specific technical errors by simultaneously inferring the cells’ phylogeny and SNVs. SCIΦ employs a Markov chain Monte Carlo (MCMC) algorithm to sample the joint posterior distribution of SNVs and the phylogenetic tree of the single cells and reports the tree(s) with the best posterior probability and the corresponding genotypes. scVILP is formulated as an instance of Mixed Integer Linear Programming (MILP) and it aims to find maximum likelihood estimation (MLE) of the observed read counts given the underlying genotype matrix. Here, the MILP solver is restricted to proposing only the genotype matrices that satisfy three-gametes condition in order to maximize the likelihood function (see [19, 10, 7, 11, 23, 9] for more details on work related to inference under the three-gametes condition).

Although “regularizing” the mutation detection by using a tree as a guide is a promising direction [24, 6, 15, 18], applying SCIΦ and scVILP to datasets with large number of loci such as in [8, 31] is challenged by either very long running time or large memory consumption of the methods—the major issues in SCIΦ and scVILP, respectively. Indeed, scVILP runs out of memory on all of the datasets considered in our study here, except for the smallest ones, which is why we do not report on the performance of scVILP. To address this challenge, we developed Phylovar, a likelihood-based method for phylogeny-aware inference of SNVs from scDNAseq datasets consisting of a large number of loci. To simplify likelihood calculations for large-scale data, we assume that mutations occur following an infinite-sites assumption (ISA) [24, 4, 13]. Using this model, Phylovar finds the tree topology and the placement of mutations on ancestral single-cells that maximize the likelihood of the erroneous observed read counts given the genotypes. Utilizing a vectorized formulation for likelihood calculations, Phylovar benefits from the vectorized operations in matrix manipulation packages such as `NumPy` [12] to scale up to many loci. We compared the SNV calling accuracy, memory consumption, and running time of Phylovar against those of the existing methods, Monovar and SCIΦ, through a simulation study. We found that Phylovar outperforms SCIΦ in terms of running time with the same accuracy, while being more accurate than Monovar. Furthermore, we applied our method to two biological datasets: a triple-negative breast cancer (TNBC) dataset [31] consisting of 32 single cells and 3375 candidate loci, as well as the dataset from [8] containing 16 normal human neuron cells and 2,489,545 candidate loci. For the TNBC data, Phylovar inferred 652 SNVs with “high” or “moderate” functional impact, out of which 550 (84%) were also supported by bulk sequencing. For the neuron cells, Phylovar identified 5745 SNVs with non-synonymous effects some of which were related to neurodegenerative diseases. To the best of our knowledge, Phylovar is the first scDNAseq SNV caller that can utilize the underlying tree structure even when the dataset contains millions of genomic loci.

## Methods

The input to Phylovar consists of the reference and variant count matrices, denoted by **R** = (*r*_*ij*_) ∈ ℕ_0_^*N* ×*M*^ and **V** = (*v*_*ij*_) ∈ ℕ_0_^*N* ×*M*^, where *N* and *M* represent the number of single cells and candidate loci, respectively. Each entry in **R** and **V** represents the number of reference and variant counts, respectively, at cell *i* and site *j*. These count matrices are obtained from an input file in mpileup format. Here, candidate loci are defined as the genomic loci with a significant number of variant reads. This significance is measured by a statistic test. Note that these loci may not necessarily contain SNVs since the variant reads might be artifacts of scDNAseq technical errors. In all experiments reported below, we used SCIΦ’s likelihood ratio test described in [24] to identify candidate loci for the analyses. If the total read coverage at a cell and a candidate site is less than *λ*, the corresponding entry is treated as missing data. We used *λ* = 1 in practice.

### Single-cell genotype error model

Our genotype model considers bi-allelic genotype with 0 and 1 representing the absence and presence of a mutation, respectively. We differentiate true genotypes from those being subject to scDNAseq errors that propagate from WGA to the sequencing library — called library genotypes. Let **G** = (*g*_*ij*_) ∈ {0, 1}^*N* ×*M*^ be the binary matrix containing the true genotypes where *g*_*ij*_ represents the true genotype at cell *i* and locus *j*. Similarly, we denote the library genotype matrix by **Ψ** = (*ψ*_*ij*_) ∈ {0, 1}^*N* ×*M*^. We assume library preparation process introduces FP and FN errors into the data resulting in difference between true genotypes and their corresponding library genotypes. Let *α* and *β* denote the FP and FN error rates, respectively. Then, the probability of the library genotype given true genotype and error rates are given by the following error model adopted from SiFit [34] and SiCloneFit [33]:

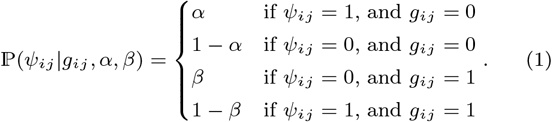

### Single-cell read count model

For the convenience of notation, let *c*_*ij*_ = *r*_*ij*_ + *v*_*ij*_ denote the total read coverage at cell *i* and locus *j*. We assume that variant read counts follow a binomial distribution whose success probability depends on the value of the library genotype:

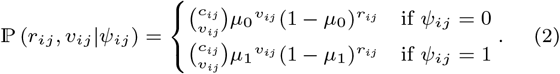

The variables *μ*_0_ and *μ*_1_ are the success probabilities associated with reference and alternate alleles, respectively. In practice, we set *μ*_0_ to 0.001, which is at the same order of magnitude for the error rate in different Illumina sequencing platforms [26]. We used 0.5 for the value of *μ*_1_, which is the mean of variant read counts for a heterozygous mutation.

### Tree model

Our tree model consists of two components: a binary tree topology *T* = (*V, E*)—where *V* denotes the set of nodes, and *E* is the set of edges—and a mutation placement for each genomic locus *j*, ℳ _*j*_ ∈ {0, 1}^2*N* −1^. The latter is a binary vector of length 2*N* −1 containing binary elements for each leaf or internal node in *V*. We take ℳ_*j*_ [*q*] = 1 to denote that a mutation occurred at node *q* during the evolutionary history of locus *j*. In our model, we assume mutations evolve following the ISA. According to this model, at most one element in ℳ _*j*_ is allowed to be 1. This vector requires each node to have an index from {1, · · ·, 2*N* − 1}. For simplicity, we map indices {1, · · ·, *N*} to the leaves/single cells and use the same mapping for the single cells in all tree topologies.

### Log-likelihood function

Assuming independence across sites/loci, the log-likelihood function of read counts given true genotypes **G**, error rates (*α, β*), underlying tree topology *T*, and mutation placements for all loci (denoted by ℳ) are:

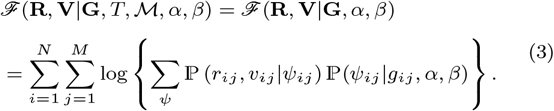

Note that the above likelihood is based on **G** rather than *T* and ℳ directly, as **G** is derived from *T* and ℳ. Therefore we can drop *T* and ℳ from Eq. (3). It can be shown that after marginalizing out *ψ*’s, log ℙ (*r*_*ij*_, *v*_*ij*_ |*g*_*ij*_, *α, β*) = log{ Σ_*ψ*_ *ℙ* (*r*_*ij*_, *v*_*ij*_ |*ψ*_*ij*_) ℙ (*ψ*_*ij*_ |*g*_*ij*_, *α, β*)} can be simplified as follows:

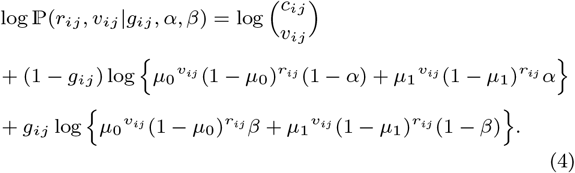

Here, the log-likelihood values of the missing data are assumed to be 0. The MLE solution is obtained as

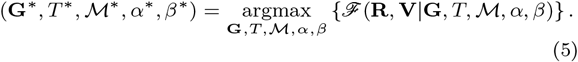

### Hill-climbing search algorithm

Phylovar infers the underlying phylogeny of single-cells and their genotypes simultaneously in a hill-climbing fashion. At each step, the log-likelihood function is evaluated and updated by proposing one of the underlying parameters including the tree, mutation placements, and error rates. We start the search by reconstructing an initial tree topology. To obtain this tree, first, we create the matrix of initial genotypes, **G**^(0)^, as follows:

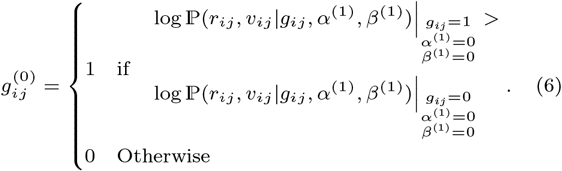

Here, (*α*^(1)^, *β*^(1)^) = (0, 0) are the initial estimates of the error rates. Given **G**^(0)^, we calculate the pairwise Hamming distances between the single cells and build an initial tree topology, *T* ^(1)^, using the neighbor-joining algorithm [22]. Given the proposed parameters (*T* ^(1)^, *α*^(1)^, *β*^(1)^), the mutation placement with highest log-likelihood for each site *j*—denoted by 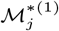—is determined yielding the genotype matrix at first iteration, **G**^(1)^ and the first log-likelihood value *ℱ* (**R, V**|**G**^(1)^, *T* ^(1)^, ℳ^*(1)^, *α*^(1)^, *β*^(1)^). At each iteration *t* ≥ 2, either new error rates are estimated or a new tree is proposed by performing tree rearrangement techniques including *subtree pruning and re-grafting* (SPR), *nearest-neighbor interchange* (NNI), and swapping two random leaves. The proposed parameters are accepted if the new log-likelihood value is greater than or equal to the log-likelihood in the previous iteration. In case of stochastic hill-climbing, the acceptance probability of the newly proposed log-likelihood value is:

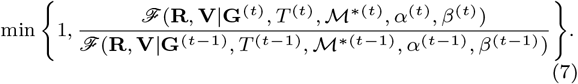

The search procedure terminates when the log-likelihood does not improve after a user-specified number of iterations or when it reaches the maximum number of iterations.

### Proposing new error rates

The new error rates at iteration *t* are calculated using the following equations from the entries of **G**^(*t*−1)^ and **G**^(0)^:

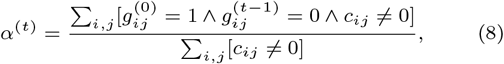

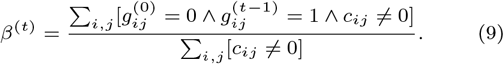

Here, the number of 0 entries in **G**^(0)^ that were “corrected” to 1 in **G**^(*t*−1)^ provides a measure of what a more realistic *α* would be through the hill-climbing trajectory. The same rationale applies to proposing a new value of *β*.

### Finding the best mutation placement

Given a topology *T* ^(*t*)^ and a site *j*, each possible mutation placement on *T* ^(*t*)^ yields a unique genotype configuration at the level of single cells. We seek the mutation placement with the highest log-likelihood. Let 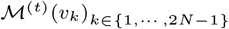 denote the mutation placement when the *k*^th^ node is mutated. As a special case, let 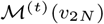 represent the absence of mutation. Because the set of all possible mutation placements is the same for all the sites, we dropped the index *j* from these two notations. To summarize the effect of all possible ISA mutation placements on genotype configurations, we define 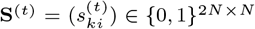 whose *k*^th^ row, 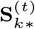, represents the genotype configuration corresponding to ℳ^(*t*)^(*v*_*k*_). We formally define the mapping from mutation placements to genotypes, denoted by Φ, as follows:

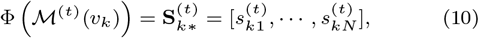

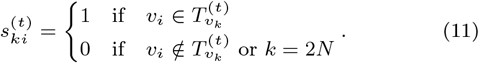

where *i* ∈ {1, · · ·, *N*} and *k* ∈ {1, · · ·, 2*N*}. Here, 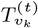 denotes the subtree rooted at node *v*_*k*_. Note that the mapping Φ is one-to-one, so we can use its inverse to retrieve the genotypes given a mutation placement. In addition to **S**^(*t*)^, we define two other matrices that store the log-likelihood values from Eq. (4), one assuming all genotypes are 0, called matrix of zero-allele likelihoods, **Z**^(*t*)^, and the other assuming all genotypes are 1, called matrix of one-allele likelihoods, **O**^(*t*)^. Formally, we define the matrices 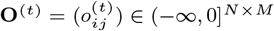 and 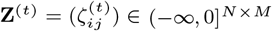 as follows:

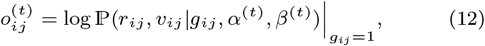

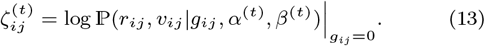

It can be shown that the following matrix multiplication results in a matrix 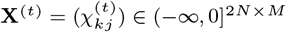 whose each element 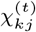 is equal to the log-likelihood value of ℳ^(*t*)^(*v*_*k*_) at site *j*:

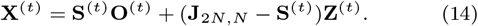

Here, **J**_2*N,N*_ is matrix of all-ones. The best mutation placement at site 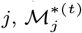, is associated with the highest value in the *j*^th^ column of **X**^(*t*)^:

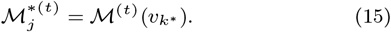

The index corresponding to the highest value is denoted by *k*^*^:

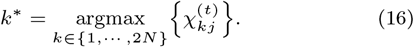

Using Φ^−1^, we can determine the best genotype configuration at site *j* which constitutes the *j*^th^ column of **G**^(*t*)^ using

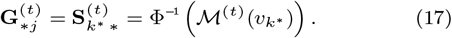

Finally, **G**^(*t*)^ is the concatenation of best genotypes configurations at all sites:

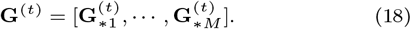

## Results and Discussion

### Simulation study

We first compared the computational efficiency and SNV calling accuracy of Phylovar, SCIΦ, and Monovar using synthetic datasets simulated under five scenarios: 1) varying the number of mutations, 2) varying the number of cells, 3) varying the ADO rates, 4) copy number effect, and 5) violation of ISA. We simulated the datasets using the simulator introduced in [24]. In the first scenario, we investigated how Phylovar performs compared to the other methods when increasing the number of mutations dramatically to a large extent.

We simulated datasets containing 16 single cells with 1000, 10^4^, and 10^5^ mutations. For each mutation value, ten datasets were generated. Phylovar’s accuracy was comparable to that of SCIΦ in terms of F1 measure, while both SCIΦ and Phylovar were more accurate than Monovar because of accounting for evolutionary history (Fig. 1a). Fig. 1f shows that the running time of each method increased with the number of mutations. For the largest dataset with 10^5^ mutations, Phylovar was approximately three orders of magnitude faster than SCIΦ.

**Fig. 1:**
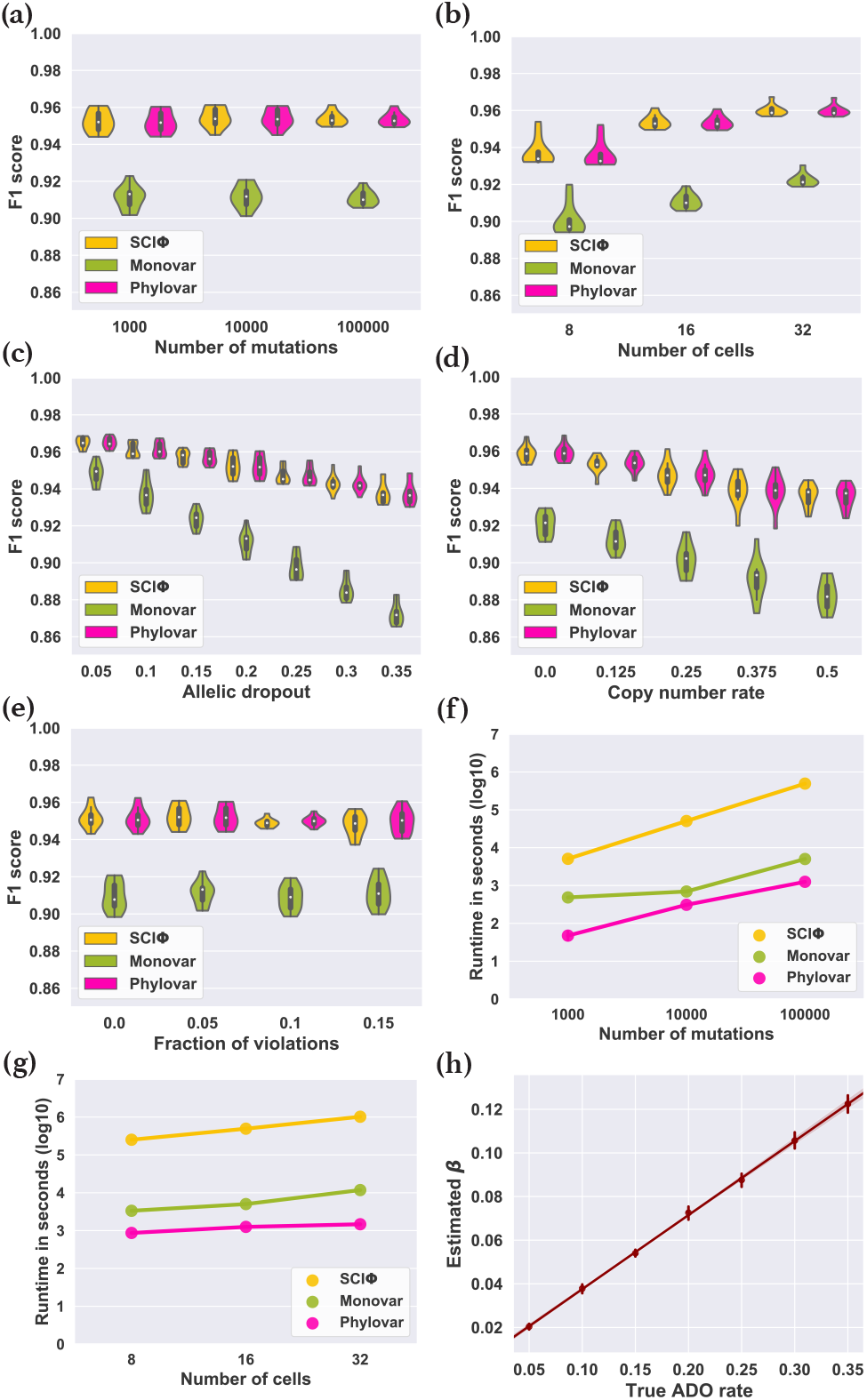
Summary statistics of different benchmarking experiments. **(a-e)** F1 accuracy of the methods from simulated data with different number of mutations **(a)**, number of cells **(b)**, ADO rate **(c)**, copy number rate **(d)**, and fraction of ISA violations **(e). (f-g)** Runtime of the methods on simulated data with varying number of mutations **(f)** and varying number of cells **(g). (h)** Linear regression between estimated false-negative error rates (*β*’s) and actual ADO rates used for simulated data.

In the second scenario, we sought to answer how the methods’ performances depend on the number of cells. Here, we fixed the number of mutations at 10^5^ and varied the number of cells, *N* ∈ {8, 16, 32}. For each setting, we generated ten datasets. The F1 accuracy scores of all methods improved as the number of cells increased (Fig. 1b). Again, Phylovar’s accuracy was comparable to that of SCIΦ while it outperformed Monovar. We observed that increasing the number of single cells improved the accuracy of all methods more than increasing the number of mutations. As demonstrated in Fig. 1g, similar to the first scenario, the running time of Phylovar was almost three orders of magnitude less than that of SCIΦ.

In the third scenario, the ADO rate was varied while cells and mutations were fixed at 16 and 1000. We selected ADO rates from the values {0, 0.05, 0.1, 0.15, 0.2, 0.25, 0.3, 0.35}, and generated ten datasets for each ADO value. Fig. 1c shows that both SCIΦ and Phylovar were more robust to high ADO rates than Monovar due to the utilization of the underlying single-cell phylogeny.

Since Phylovar can estimate the false-negative error rates (denoted by *β*), we measured the correlation between the ADO rates used for generating the simulated datasets and the estimated *β*’s. As demonstrated in Fig. 1h, these two values were highly correlated (the Pearson correlation coefficient was 0.991). It is worth noting that based on the linear regression line, the estimated *β* was almost half of the true ADO, pointing to the difference between the dropout mechanism in the simulator and our definition of *β* (see Methods). Given an ADO rate *μ*, the simulator chooses *μ* fraction of the mutations. It changes 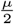 of them into reference genotype, and 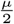 of them into homozygous mutations; the *β* in our model indicates the probability of a mutation becoming reference genotype implying 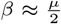, which we can observe in Fig. 1h.

Since Phylovar assumes the read counts are originated from diploid strands, in the fourth scenario, additional wild type alleles were introduced to the read counts to imitate the effect of copy number changes. The simulator randomly selects a fraction of mutated loci (named copy number rate), and chooses *c* extra copies for each loci with probability 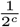 [24]. We increased the copy number rate from 0 to 0.5 with step size 0.125. For each value, we generated ten datasets containing 16 cells and 1000 mutations. Fig. 1d shows that the SNV calling accuracy of the methods decreased as more mutated loci were subject to copy number changes.

In the fifth scenario, we were interested in observing how violations of ISA affect the SNV calling accuracy of the methods. Given a fraction of mutations, the simulator randomly selects half of them to recur in different branches, and the rest of them to be lost in the same subtree. We increased the fraction of mutations subject to ISA violations from 0 to 0.15 with 0.05 step size. For each value we generated ten datasets with 16 cells and 1000 mutations. Fig. 1e shows that all three methods had a stable performance as the fraction of ISA violations increased. This observation suggests that even though the phylogenies inferred by SCIΦ and Phylovar might be inaccurate due to the presence of violations of their evolutionary model, the effect of such violations on mutation inference is negligible.

### Application to real data

We applied Phylovar on two human scDNAseq datasets. The first dataset consists of single-cell whole-exome sequencing (scWES) samples from a triple-negative breast cancer (TNBC) patient [31]. Since the population sequencing data from bulk tumor and matched normal tissue are available for the TNBC dataset, the number of mutations shared by scWES and bulk data provides us a metric for measuring the accuracy of our approach. The TNBC dataset consists of 16 diploid cells, eight hyperdiploid/aneuploid cells, and eight hypodiploid cells [31]. Given the control normal cells, SCIΦ’s likelihood ratio test identifies the loci likely to contain somatic mutations. Applying this statistic test on the input mpileup file resulted in 3375 candidate loci on which we applied Phylovar. Phylovar was run with ten parallel hill-climbing chains, each for 100,000 iterations on a pool of five CPU’s, each with 48 cores (AMD EPYC 7642) on a node with 192 GB RAM. The total runtime was 91 minutes. Phylovar inferred an 18.21% false-negative error rate and a 1.03% false-positive error rate from TNBC data. We ran SCIΦ and Monovar with default parameters; SCIΦ and Monovar terminated after 10 hours and 144 minutes, respectively. Fig. 2 shows the three methods’ mutation calls on TNBC data from the overlapping sites as well as the initial genotype matrix at the first iteration of our hill-climbing search. We performed hierarchical clustering with Ward’s minimum variance method implemented in Python’s `SciPy` package [29] on the genotype matrix for better visualization. We observed concordance between the calls from Phylovar and SCIΦ while Monovar’s calls are noisy and resemble Phylovar’s initial genotypes.

**Fig. 2:**
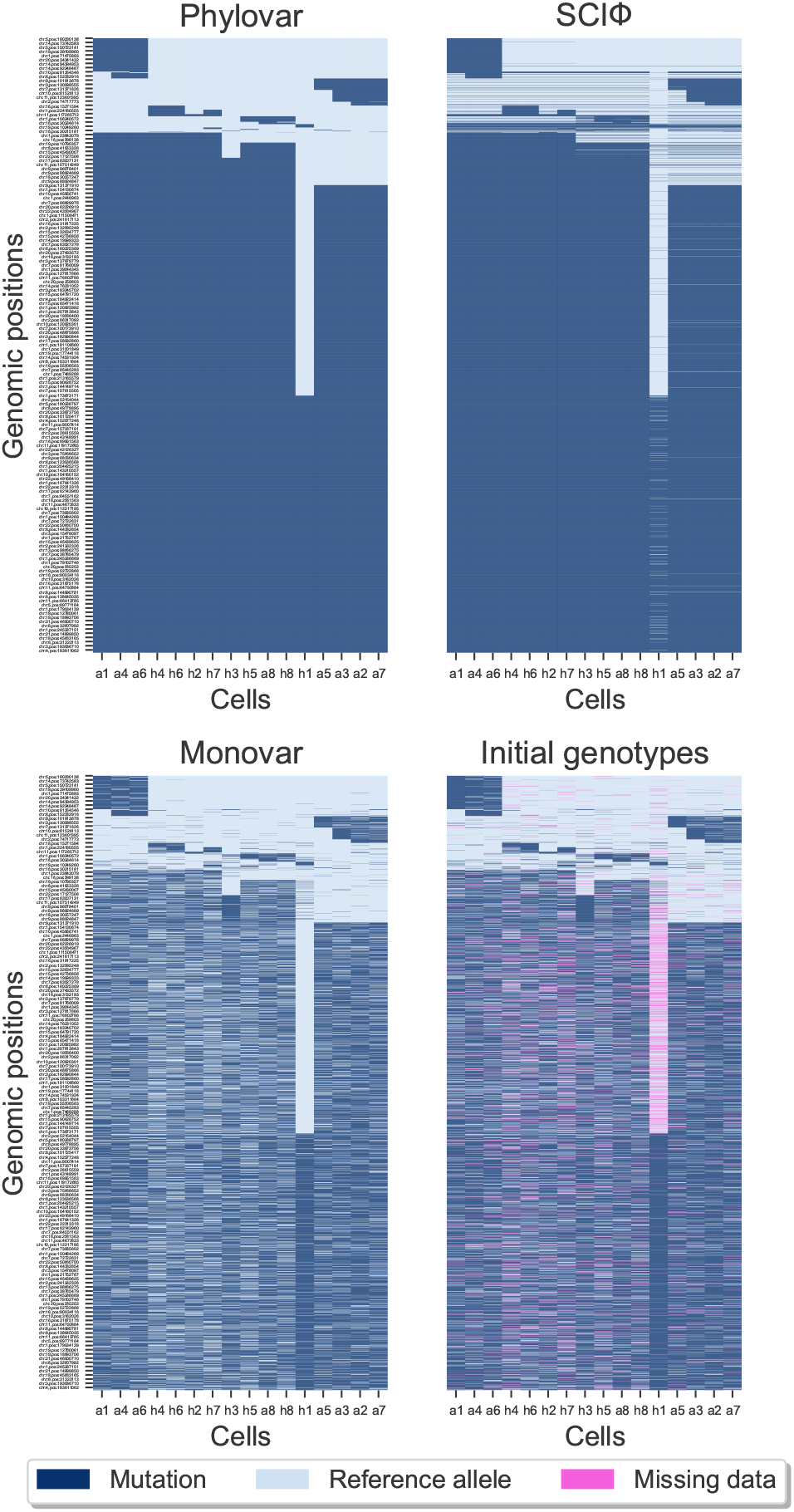
Clustered heatmaps of mutation calls by different approaches performed on the TNBC dataset. Here, rows and columns represent the genomic loci and the single-cells, respectively. The pixels show mutation calls (dark blue), reference alleles (light blue), and missing data (pink). The initial genotypes are the initial estimates of genotypes considering no error rates and no underlying phylogeny at the starting step of Phylovar’s search algorithm.

To annotate the mutations, we applied snpEff [1] on the SNVs detected by Phylovar. Out of 3375 candidate loci, 652 loci contained SNVs with “high” or “moderate” functional effects (see [2] for details on the types of variants’ effects and their descriptions). Then, we ran HaplotypeCaller (GATK version 4.2.0.0) for mutation calling on the bulk tumor and normal samples. Among 652 SNVs in single cells, 550 (84%) mutations were found in bulk data (Fig. 3).

**Fig. 3:**
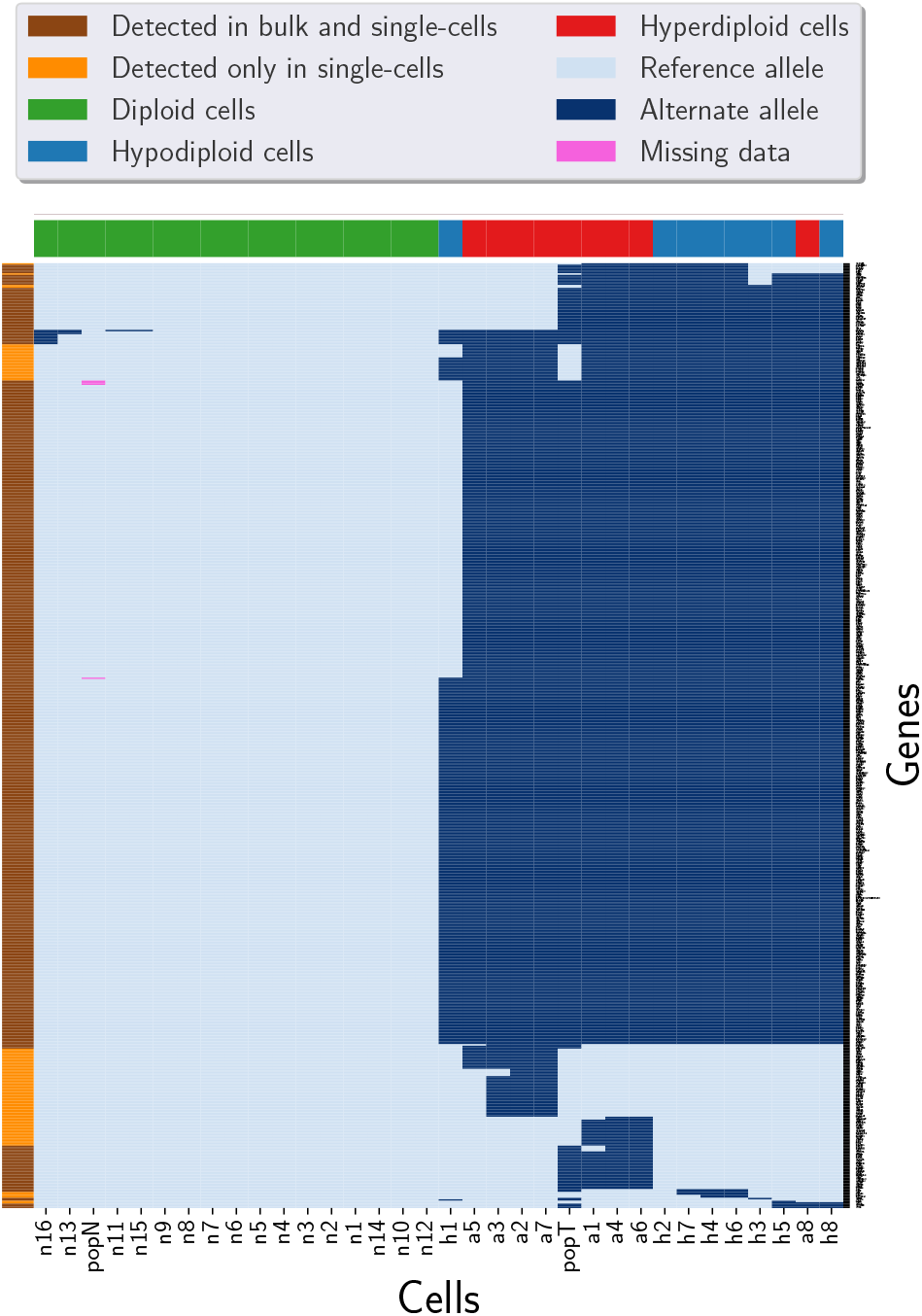
Clustered heatmap of the mutations detected from TNBC data and population sequencing data of tumor and matched normal tissues. The pixels show mutation calls (dark blue), reference alleles (light blue), and missing data (pink). Columns (cells) are colored according to the ploidy of the cells. The colors of the rows (genes) indicate whether the SNV was found in bulk data or not. Here, `popT` and `popN` are the tumor and normal population sequencing samples, respectively. Out of 3375 candidate loci, 652 loci contained SNVs with high or moderate functional effects in the single-cell data among which 550 mutations were found in bulk data as well, which is 84% of the single-cell calls.

The second biological data consists of 16 neuron cells on which scWGS was performed to study somatic mutations in human brain development [8]. Applying SCIΦ’s statistic test on the input mpileup identified 2,489,545 candidate loci. We ran Phylovar with five parallel hill-climbing chains, each for 50,000 iterations on five CPUs with 192 GB RAM. Phylovar finished the process after 17 hours and 45 minutes. To compare our results with other methods, we ran Monovar and SCIΦ with default parameters. Monovar processed the data in 10 hours and 26 minutes, while SCIΦ was still running after ten days. Phylovar’s inferred false-positive and false-negative error rates were 75.22%, and 1.17%, respectively. snpEff identified 5745 non-synonymous SNVs among Phylovar’s mutation calls. Fig. 4 shows hierarchical clustering on the genotypes of Phylovar, Monovar, and the initial genotypes at sites with non-synonymous SNVs. We observed similarities between Monovar’s calls and the initial genotypes. By comparing the panel of Phylovar in Fig. 4 with the other panels, one can see the sparse regions of mutations in panels of Monovar and the initial genotype matrix that are inferred as reference alleles by Phylovar.

**Fig. 4:**
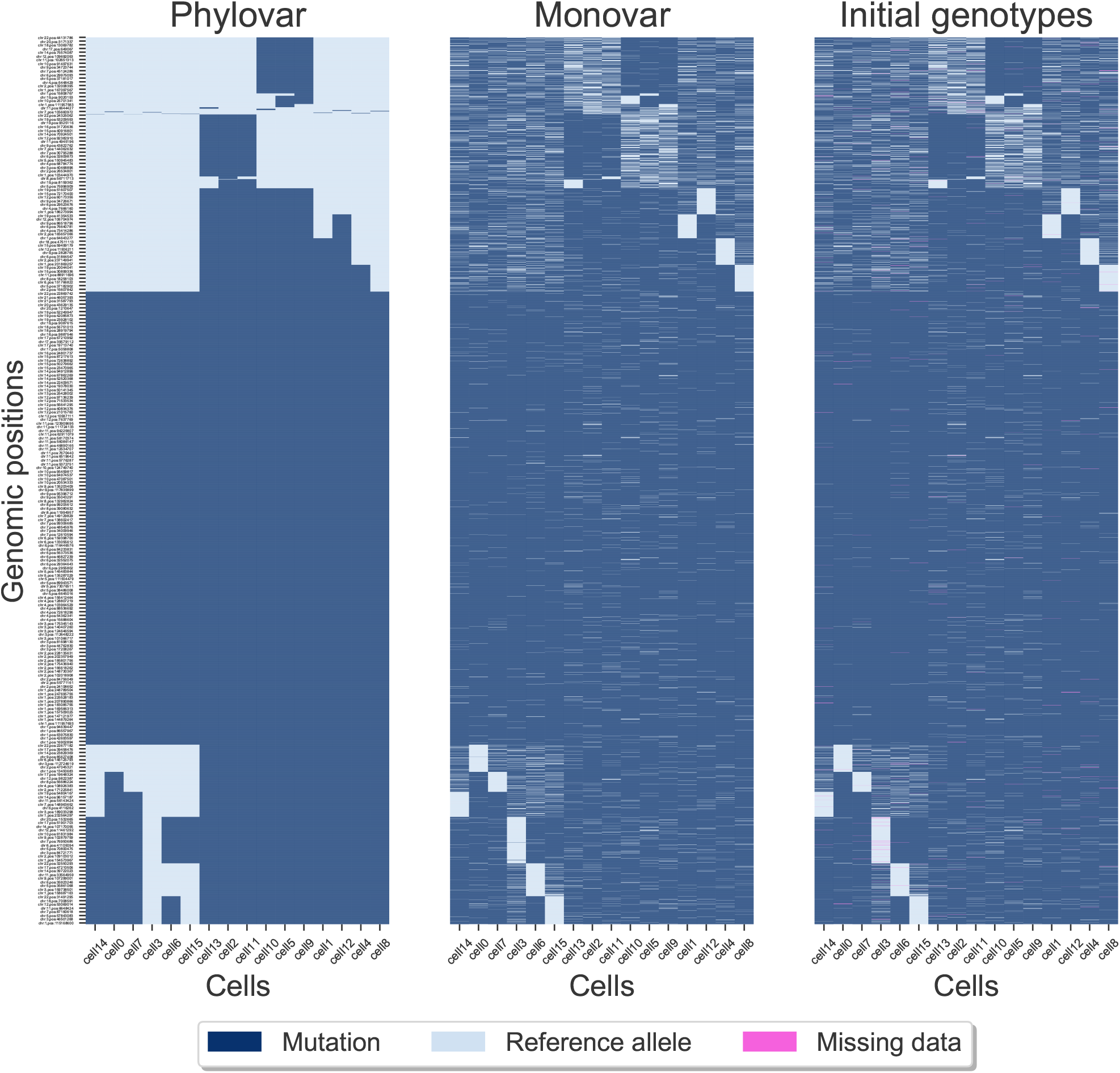
Clustered heatmaps of mutation calls by different approaches performed on neuron cells. Rows and columns represent the genomic loci and the single-cells, respectively. The pixels show mutation calls (dark blue), reference alleles (light blue), and missing data (pink). The initial genotypes are the initial estimates of genotypes considering no error rates and no underlying phylogeny at the starting step of Phylovar’s search algorithm.

Further, we investigated the genes likely related to neurodegenerative diseases by comparing our findings with the genes reported in [32]. Wei et al. [32] studied somatic mutations in 1461 control and diseased human brains with different neurodegenerative disorders. Among the genes inferred by Phylovar to harbor non-synonymous mutations, 12 genes were reported in [32]. We observed that the genes *MUC16* and *MLIP* were frequently mutated in different regions; also, non-synonymous SNVs were observed within *KRT33A* and *SEMA5B* in [32] from patients with Creutzfeldt-Jakob and Alzheimer diseases, respectively. The presence of these non-synonymous mutations in both diseased and normal samples implies the high mutability of these genes even in a healthy individual.

## Conclusions

The rapid growth of SCS technologies poses computational challenges due to the increasing number of cells and sites sequenced per genome [17]. In this work, we focused on addressing the computational challenge associated with the breadth of genomic sites in scDNAseq data. Here, we introduced Phylovar, a scalable MLE method for phylogeny-guided inference SNVs from single-cell DNA sequencing data suitable for scWGS and scWES data with an extensive number of loci. We introduced a novel vectorized formula for likelihood calculation, making Phylovar scalable to hundreds of thousands, even millions of loci.

We assessed Phylovar’s performance against state-of-the-art variant callers SCIΦ [24] and Monovar [36], through simulated benchmarks. Phylovar outperforms SCIΦ in terms of running time while being more accurate than Monovar in different simulation scenarios. We also applied Phylovar to two real biological datasets. For a TNBC dataset with 32 single cells and 3375 candidate loci, Phylovar identified SNVs with functional impact among which 84% were supported by bulk sequencing data. Phylovar was also more accurate than Monovar and 6.5x faster than that of SCIΦ. For a larger dataset containing 16 normal human neuron cells and approximately 2.5 million candidate loci, Phylovar identified 5745 non-synonymous SNVs some of which were related to neurodegenerative diseases. Interestingly, Phylovar detected 75.22% false-positive, and 1.17% false-negative error rate for this dataset. The neuron cells data was particularly challenging due to large number of sites. For this data, SCIΦ failed to converge even after ten days of running while Phylovar terminated after less than 18 hours.

Phylovar makes it possible to analyze datasets with large number of loci within reasonable time and memory requirements, thus adding to the growing toolbox for analyzing scDNAseq data. As a direction for future research, we will explore deviations from the simplified ISA model and investigate the feasibility of applying more general finite-sites models (FSM) [34, 33] to datasets with many loci. As scDNAseq technologies advance, the sequencing cost per cell decreases [16, 28]. Consequently, we expect more scWGS and scWES datasets to emerge in the future, requiring methods such as Phylovar that can perform scalable variant calling on datasets with millions of loci.

## Acknowledgements

We would like to thank Dr. Nicholas Navin and Min Hu for providing access to the TNBC population sequencing data. Also, we thank Prof. Dan Gusfield and Dr. Robert Gysel for their insight on the concept of Perfect Phylogeny and helping us with testing their tools.

This work was supported in part by the National Science Foundation, grants IIS-1812822 and IIS-2106837 (L.N.).

## References

1. P. Cingolani, A. Platts, M. Coon, T. Nguyen, L. Wang, S.J. Land, X. Lu, and D.M. Ruden. A program for annotating and predicting the effects of single nucleotide polymorphisms, snpeff: Snps in the genome of drosophila melanogaster strain w1118; iso-2; iso-3. Fly, 6(2):80–92, 2012.

2. Pablo Cingolani. Input & output files. https://pcingola.github.io/SnpEff/se_inputoutput/.

3. Frank B. Dean, Seiyu Hosono, Linhua Fang, Xiaohong Wu, A. Fawad Faruqi, Patricia Bray-Ward, Zhenyu Sun, Qiuling Zong, Yuefen Du, Jing Du, Mark Driscoll, Wanmin Song, Stephen F. Kingsmore, Michael Egholm, and Roger S. Lasken. Comprehensive human genome amplification using multiple displacement amplification. PNAS, 99(8):5261– 5266, 2002.

4. Amit G. Deshwar, Shankar Vembu, Christina K. Yung, Gun Ho Jang, Lincoln Stein, and Quaid Morris. PhyloWGS: Reconstructing subclonal composition and evolution from whole-genome sequencing of tumors. Genome Biology, 16(1):35, 2015.

5. Xiao Dong, Lei Zhang, Brandon Milholland, Moonsook Lee, Alexander Y Maslov, Tao Wang, and Jan Vijg. Accurate identification of single-nucleotide variants in whole-genome-amplified single cells. Nature Methods, 14(5):491–493, 2017.

6. Mohammadamin Edrisi, Hamim Zafar, and Luay Nakhleh.A Combinatorial Approach for Single-cell Variant Detection via Phylogenetic Inference. 19th International Workshop on Algorithms in Bioinformatics (WABI 2019), 143:1–13, 2019.

7. G. Estabrook, C. Johnson, and F. McMorris. A mathematical formulation for the analysis of cladistic chatracter compatibility. Mathematical Biosciences, 29:181–187, 1976.

8. Gilad D Evrony, Eunjung Lee, Bhaven K Mehta, Yuval Benjamini, Robert M Johnson, Xuyu Cai, Lixing Yang, Psalm Haseley, Hillel S Lehmann, Peter J Park, and Christopher A Walsh. Cell lineage analysis in human brain using endogenous retroelements. Neuron, 85(1):49–59, 01 2015.

9. David Fernández-Baca. Steiner Trees in Industry, volume 11, chapter The Perfect Phylogeny Problem, pages 203–234. Springer, Boston, MA, US, 01 2001.

10. D. Gusfield. Efficient algorithm for inferring evolutionary trees. Networks, 21(1):19–28, 1991.

11. D. Gusfield. Algorithms on Strings, Trees and Sequences. Cambridge University Press, 1997. Cambridge, UK.

12. Charles R. Harris, K. Jarrod Millman, Stéfan J van der Walt, Ralf Gommers, Pauli Virtanen, David Cournapeau, Eric Wieser, Julian Taylor, Sebastian Berg, Nathaniel J. Smith, Robert Kern, Matti Picus, Stephan Hoyer, Marten H. van Kerkwijk, Matthew Brett, Allan Haldane, Jaime Fernández del Río, Mark Wiebe, Pearu Peterson, Pierre Gérard-Marchant, Kevin Sheppard, Tyler Reddy, Warren Weckesser, Hameer Abbasi, Christoph Gohlke, and Travis E. Oliphant. Array programming with NumPy. Nature, 585:357—-362, 2020.

13. Katharina Jahn, Jack Kuipers, and Niko Beerenwinkel. Tree inference for single-cell data. Genome Biology, 17(1):86, 2016.

14. Yukie Kashima, Yoshitaka Sakamoto, Keiya Kaneko, Masahide Seki, Yutaka Suzuki, and Ayako Suzuki. Single-cell sequencing techniques from individual to multiomics analyses. Experimental & Molecular Medicine, 52(9):1419– 1427, 2020.

15. Jack Kuipers, Mustafa Anıl Tuncel, Pedro Ferreira, Katharina Jahn, and Niko Beerenwinkel. Single-cell copy number calling and event history reconstruction. bioRxiv, 2020.

16. Bora Lim, Yiyun Lin, and Nicholas Navin. Advancing Cancer Research and Medicine with Single-Cell Genomics. Cancer Cell, 37(4):456–470, 2020.

17. David Lähnemann, Johannes Köster, Ewa Szczurek, Davis J. McCarthy, Stephanie C. Hicks, Mark D. Robinson, Catalina A. Vallejos, Kieran R. Campbell, Niko Beerenwinkel, Ahmed Mahfouz, Luca Pinello, Pavel Skums, Alexandros Stamatakis, Camille Stephan-Otto Attolini, Samuel Aparicio, Jasmijn Baaijens, Marleen Balvert, Buys de Barbanson, Antonio Cappuccio, Giacomo Corleone, Bas E. Dutilh, Maria Florescu, Victor Guryev, Rens Holmer, Katharina Jahn, Thamar Jessurun Lobo, Emma M. Keizer, Indu Khatri, Szymon M. Kielbasa, Jan O. Korbel, Alexey M. Kozlov, Tzu-Hao Kuo, Boudewijn P. F. Lelieveldt, Ion I. Mandoiu, John C. Marioni, Tobias Marschall, Felix Mölder, Amir Niknejad, Lukasz Raczkowski, Marcel Reinders, Jeroen de Ridder, Antoine-Emmanuel Saliba, Antonios Somarakis, Oliver Stegle, Fabian J. Theis, Huan Yang, Alex Zelikovsky, Alice C. McHardy, Benjamin J. Raphael, Sohrab P. Shah, and Alexander Schönhuth. Eleven grand challenges in single-cell data science. Genome Biology, 21(1):31, 2020.

18. Magda Markowska, Tomasz Cakala, Blazej Miasojedow, Dilafruz Juraeva, Johanna Mazur, Edith Ross, Eike Staub, and Ewa Szczurek. Conet: Copy number event tree model of evolutionary tumor history for single-cell data. bioRxiv, 2021.

19. C. Meacham. Numerical Taxonomy. NATO ASI Series (Series G: Ecological Sciences), volume 1, chapter Theoretical and computational considerations of the compatibility of qualitative taxonomic characters. Springer, Berlin, Germany, 1983.

20. Nicholas Navin, Jude Kendall, Jennifer Troge, Peter Andrews, Linda Rodgers, Jeanne McIndoo, Kerry Cook, Asya Stepansky, Dan Levy, Diane Esposito, Lakshmi Muthuswamy, Alex Krasnitz, W. Richard McCombie, James Hicks, and Michael Wigler. Tumour evolution inferred by Inference of single-nucleotide variations from single-cell DNA 9 single-cell sequencing. Nature, 472(7341):90–94, 2011.

21. Nicholas E. Navin. Cancer genomics: one cell at a time. Genome Biology, 15(8):452, 2014.

22. N Saitou and M Nei. The neighbor-joining method: a new method for reconstructing phylogenetic trees. Molecular Biology and Evolution, 4(4):406–425, 07 1987.

23. C. Semple and M. Steel. Phylogenetics. Oxford University Press, 2003. UK.

24. Jochen Singer, Jack Kuipers, Katharina Jahn, and Niko Beerenwinkel. Single-cell mutation identification via phylogenetic inference. Nature Communications, 9(1):5144, 2018.

25. Claudia Spits, Cédric Le Caignec, Martine De Rycke, Lindsey Van Haute, André Van Steirteghem, Inge Liebaers, and Karen Sermon. Whole-genome multiple displacement amplification from single cells. Nature Protocols, 1(4):1965– 1970, 2006.

26. Nicholas Stoler and Anton Nekrutenko. Sequencing error profiles of Illumina sequencing instruments. NAR Genomics and Bioinformatics, 3(1), 03 2021.

27. Fuchou Tang, Catalin Barbacioru, Yangzhou Wang, Ellen Nordman, Clarence Lee, Nanlan Xu, Xiaohui Wang, John Bodeau, Brian B Tuch, Asim Siddiqui, Kaiqin Lao, and M Azim Surani. mRNA-Seq whole-transcriptome analysis of a single cell. Nature Methods, 6(5):377–382, 2009.

28. Xiaoning Tang, Yongmei Huang, Jinli Lei, Hui Luo, and Xiao Zhu. The single-cell sequencing: new developments and medical applications. Cell & Bioscience, 9(1):53, 2019.

29. Pauli Virtanen, Ralf Gommers, Travis E. Oliphant, Matt Haberland, Tyler Reddy, David Cournapeau, Evgeni Burovski, Pearu Peterson, Warren Weckesser, Jonathan Bright, Stéfan J. van der Walt, Matthew Brett, Joshua Wilson, K. Jarrod Millman, Nikolay Mayorov, Andrew R. J. Nelson, Eric Jones, Robert Kern, Eric Larson, CJ Carey, Ilhan Polat, Yu Feng, Eric W. Moore, Jake VanderPlas, Denis Laxalde, Josef Perktold, Robert Cimrman, Ian Henriksen, E. A. Quintero, Charles R. Harris, Anne M. Archibald, Antônio H. Ribeiro, Fabian Pedregosa, Paul van Mulbregt, and SciPy 1.0 Contributors. SciPy 1.0: Fundamental Algorithms for Scientific Computing in Python. Nature Methods, 17:261–272, 2020.

30. Yong Wang and Nicholas E. Navin. Advances and Applications of Single-Cell Sequencing Technologies. Molecular Cell, 58(4):598–609, 2015.

31. Yong Wang, Jill Waters, Marco L. Leung, Anna Unruh, Whijae Roh, Xiuqing Shi, Ken Chen, Paul Scheet, Selina Vattathil, Han Liang, Asha Multani, Hong Zhang, Rui Zhao, Franziska Michor, Funda Meric-Bernstam, and Nicholas E. Navin. Clonal evolution in breast cancer revealed by single nucleus genome sequencing. Nature, 512(7513):155–160, 2014.

32. Wei Wei, Michael J. Keogh, Juvid Aryaman, Zoe Golder, Peter J. Kullar, Ian Wilson, Kevin Talbot, Martin R. Turner, Chris-Anne McKenzie, Claire Troakes, Johannes Attems, Colin Smith, Safa Al Sarraj, Chris M. Morris, Olaf Ansorge, Nick S. Jones, James W. Ironside, and Patrick F. Chinnery. Frequency and signature of somatic variants in 1461 human brain exomes. Genetics in Medicine, 21(4):904–912, 2019.

33. H. Zafar, N. Navin, K. Chen, and L. Nakhleh. SiCloneFit: Bayesian inference of population structure, genotype, and phylogeny of tumor clones from single-cell genome sequencing data. Genome Research, 29:1860–1877, 2019.

34. H. Zafar, A. Tzen, N. Navin, K. Chen, and L. Nakhleh. SiFit: inferring tumor trees from single-cell sequencing data under finite-sites models. Genome Biology, 18(1):178, 2017.

35. Hamim Zafar, Nicholas Navin, Luay Nakhleh, and Ken Chen. Computational approaches for inferring tumor evolution from single-cell genomic data. Current Opinion in Systems Biology, 7:16–25, 2018.

36. Hamim Zafar, Yong Wang, Luay Nakhleh, Nicholas Navin, and Ken Chen. Monovar: single-nucleotide variant detection in single cells. Nature Methods, 13(6):505–507, 2016.

37. Chenghang Zong, Sijia Lu, Alec Chapman, and X Xie. Genome-Wide Detection of Single-Nucleotide and Copy-Number Variations of a Single Human Cell. Science, 338:1622–1666, 12 2012.

